# More motivated but equally good: no effect of gamification on visual working memory performance

**DOI:** 10.1101/2020.01.12.903203

**Authors:** Maria Mystakidou, Ronald van den Berg

## Abstract

Gamification refers to the introduction of gaming elements such as scores and leaderboards in non-gaming contexts. While there is growing evidence that gamification has positive effects on intrinsic motivation and engagement, it is largely unknown whether these effects translate to improved cognitive performance. Here, we examine whether gamification affects performance on a visual working memory (VWM) task. In Experiment 1, we gamified a standard delayed-estimation task by introducing scores and a leveling system. On each trial, the subject’s estimation error was mapped to a score between −100 and +100 and added to their total score. Subjects started at a set size of 1 and “leveled up” to the next set size each time they had accumulated 1,500 points. Post-experiment questionnaire data confirmed that subjects who performed the gamified version of the task were more motivated than control subjects. However, we found no difference in VWM performance between these two groups, nor between below-median and above-median motivated subjects. In Experiment 2, we tested for effects of trial-by-trial manipulations of motivation on VWM performance, by varying the scoring function across trials. Three scoring functions were used, with maxima of 7, 21, and 101 points. At the beginning of each trial, the subject was informed whether the potential reward was “low”, “medium”, or “high”. Post-questionnaire data showed that subjects were more motivated on high-reward trials. However, we found no evidence for a difference in performance between the three reward levels. Our results suggest that gamification increases people’s motivation to carry out visual working memory tasks, but it does not necessarily increase their performance.

## 1. INTRODUCTION

There has recently been a surge in research on effects of “gamification”, i.e., the use of gaming elements in non-gaming contexts such as learning, education, and marketing (e.g., (Deterding, Dixon, Khaled, & Nacke, 2011; Dicheva, Dichev, Agre, & Angelova, 2015; Hamari & Lehdonvirta, 2010; Hanus & Fox, 2015; Subhash & Cudney, 2018)). Empirical work has suggested that gamification has positive effects on a variety of psychological outcomes, such as intrinsic motivation, enjoyment, engagement, and perceived competence (see Hamari et al., 2014 for a review). It is largely unknown, however, whether these effects are accompanied by improved cognitive performance. In the present study, we examine whether gamification improves people’s performance on a visual working memory (VWM) task.

A prerequisite for finding effects of gamification on VWM performance is that allocation of VWM resources must be flexible – if it is fixed, no experimental manipulation can increase or decrease VWM performance. While there has been extensive research on describing VWM limitations (Brady, Konkle, & Alvarez, 2011; Luck & Vogel, 2013; Ma, Husain, & Bays, 2014), few studies have asked why there are limitations in the first place. One possible answer to this question is that the sustained energy that is required to keep a memory alive (Fuster & Alexander, 1971) induces a metabolic cost (Attwell & Laughlin, 2001; Laughlin, 2001; Sterling & Laughlin, 2015). In the presence of such a cost, a rational system would use its resources sparingly: resources should only be invested insofar the induced cost is compensated for by expected task reward. We recently formalized this idea in a resource-rational theory of VWM (Van den Berg & Ma, 2018) and found that a model derived from this theory accounts well for earlier documented effects of set size (Bays, Catalao, & Husain, 2009; Van den Berg, Shin, Chou, George, & Ma, 2012; Wilken & Ma, 2004) and item importance (Bays, 2014; Emrich, Lockhart, & Al-Aidroos, 2017) on encoding precision. Inspired by these findings, we proposed that resource-rationality may be a general theory of how the brain allocates VWM resources and that VWM “limitations” are the result of a cost-benefit trade-off rather than a hardwired constraint on capacity. A key implication of this theory is that VWM resource allocation may be much more flexible than assumed so far.

If the amount of VWM resource utilized to a task is flexible, we may expect a relation between a subject’s level of motivation and the amount of resource they invest in a task. Therefore, we hypothesize that performance on VWM tasks can be improved by gamifying the tasks. We test this hypothesis in two experiments. Experiment 1 uses a between-subject design in which we compare VWM performance between subjects who perform a standard VWM task and subjects who perform a gamified version of that task. Experiment 2 uses a within-subject design, in which the number of points that a subject can earn varies across trials. To preview our results, we find that gamification increases motivation in both experiments, but we find no effect on VWM performance.

## 2. EXPERIMENT 1

### 2.1. Sharing of data and analysis scripts

Data and Matlab scripts related to this experiment are available at https://osf.io/gb2kd/.

### 2.2. Participants

A total of 62 participants with self-reported normal or corrected-to-normal vision were recruited using posters at various campuses of Uppsala University (Table 1). The first 40 participants were randomly divided into the first three gamified groups. The remaining 22 participants were recruited later and randomly divided into the control group and the last gamified group. The study was approved by the Regional Ethical Review Board in Uppsala and conducted according to the Declaration of Helsinki Principles. All participants signed informed consent and received a cinema voucher with a value of approximately $12 for their participation.

**Table 1.** Overview of participants. E0 refers to the estimation error (in degrees) at which the scoring function mapped to a score of 0 points (see Figure 1B).

| Condition | $E_0$ | Number of subjects | Age range | Mean age | Females |
| --- | --- | --- | --- | --- | --- |
| Gamified, difficulty 1 | 60 | 13 | 20-47 | 28.0±0.5 | 7 |
| Gamified, difficulty 2 | 45 | 13 | 19-36 | 24.4±0.4 | 9 |
| Gamified, difficulty 3 | 30 | 14 | 20-43 | 26.4±0.5 | 10 |
| Gamified, difficulty 4 | 20 | 10 | 21-38 | 25.2±0.5 | 6 |
| Control | - | 12 | 20-35 | 24.6±0.4 | 9 |

### 2.3. Stimuli and materials

Stimuli were presented on a 23” LCD screen at a resolution of 1920×1080 pixels in a dimly lighted room^1^. Subjects were seated at a distance of approximately 60 cm from the screen. Two centrally located concentric circles were used as a fixation point. Stimuli were dark gray ellipses, presented on a light gray background (Figure 1A). The ellipses had an area of 1,000 pixels^2^ and an eccentricity (elongation) of 0.95. The stimuli were presented at a centrally located, invisible circle with a radius of 220 pixels. The location of the first stimulus was drawn randomly and all other stimuli were positioned such that equal spacing was ensured between any two neighboring stimuli. The orientation of each stimulus was drawn from a uniform distribution on the circle, with the constraint that the minimum circular distance (in degrees) between any two stimuli was at least 90/*N*, where *N* indicates the set size. This constraint was intended to discourage subjects from using chunking strategies during encoding (Nassar, Helmers, & Frank, 2018). Eye movements were recorded using a Tobii 4C eye tracker.

**Figure 1.**
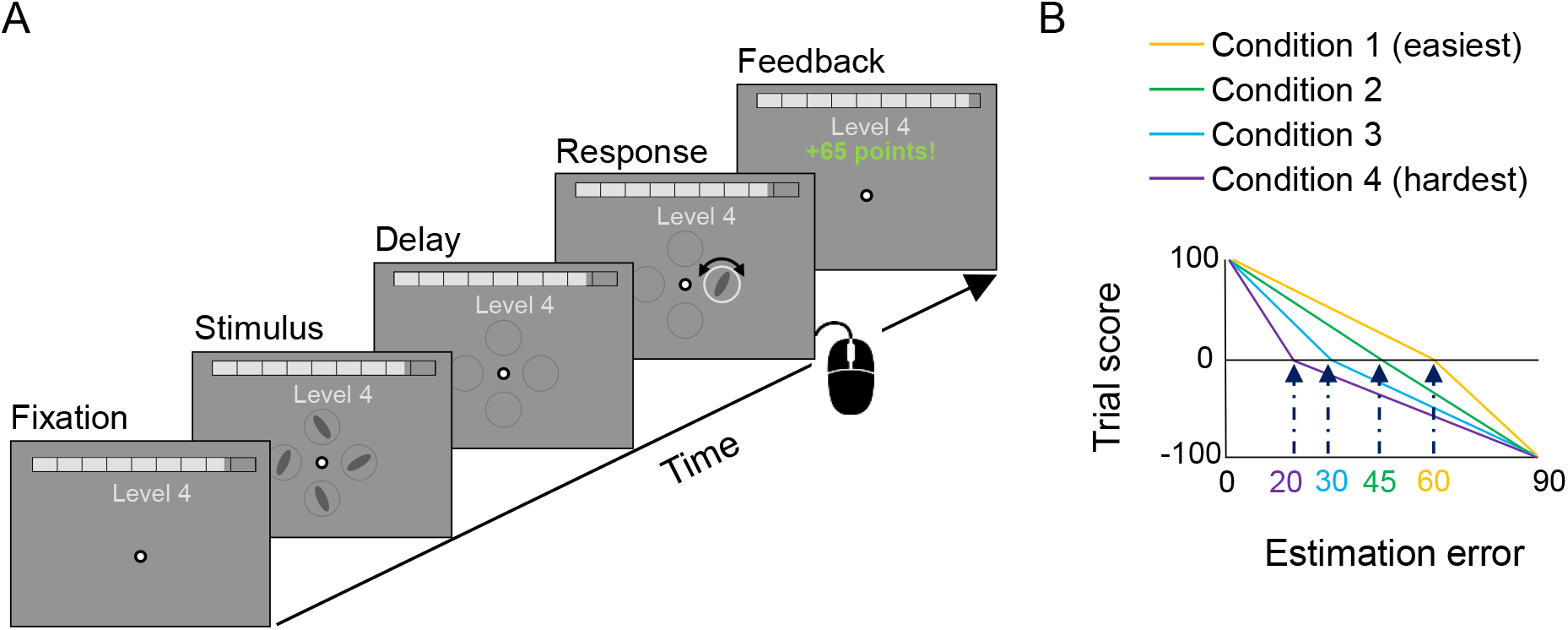
Design of Experiment 1. Schematic illustration of a single experimental trial. (B) Scoring functions used in the four gamified conditions.

### 2.4. Task

Subjects performed a delayed estimation task (Wilken & Ma, 2004) with stimulus orientation as the relevant feature. Each trial started with a central fixation mark which the subject had to fixate at for at least 600 ms. If a subject failed to fixate within 1 second, a message would appear on the screen telling the subject to “Please fixate at the central dot”. After successful fixation, the subject was presented with *N* oriented ellipses (200 ms). After a memory delay (1.5 s), the location of one ellipse was highlighted with a dashed circle. The task of the subject was to reproduce the orientation of the earlier displayed ellipse at that location. In order to respond, the subject would first click anywhere on the dashed circle to indicate their initial orientation estimate. Next, an ellipse would appear whose orientation lined up with where the subject had clicked the circle. The subject could then adjust the orientation by moving the mouse and submit the response by clicking the left mouse button. The inter-trial time was 700 ms.

#### Gamified conditions

To gamify the task, we extended it with two of the most commonly used concepts in gamification (Hamari et al., 2014): points and levels. We equated the level of the “game” with the set size, i.e., the number of elements in the stimulus display. Subjects started at level 1 and would level up to the next set size each time they had accumulated 1,500 points^2^. On each trial, they earned or lost points based on the accuracy of their response. The scoring function was a two-part linear function in which the minimum error (0°) mapped to 100 points and the maximum error (90°) to −100 points. The error magnitude that mapped to 0 points, referred to as *E*_0_, differed between the four gamified conditions (*E*_0_=20, 30, 45, 60) and determined the difficulty of the task: the higher *E*_0_, the easier it was to score points (Figure 1B). We expected that task difficulty might influence intrinsic motivation levels (e.g., lower motivation when the task was considered too easy or too hard). Throughout the experiment, a progress bar was visible at the top of the screen to indicate how far or close the subject was to reaching the next level (Figure 1A). After each trial, the subject received feedback about the number of gained or lost points and saw the progress bar grow or shrink accordingly. Visualizing subjects’ competence may be expected to further increase their intrinsic motivation for performing the task (Ryan & Deci, 2000). The gamified conditions consisted of two rounds, each starting at level 1 and lasting 30 minutes. Having two rounds allowed us to examine learning effects and to dissociate these effects from potential other effects. After each round, subjects were told on the screen that they could either “Perform an additional 20 trials to improve your score” or “Skip the extra trials”. The choice that subjects made here was used as “free choice” measure of motivation (Deci, 1971, 1972).

#### Control condition

In the control condition, no information about leveling or scores was present on the screen or in the instructions. Instead of leveling up based on performance, subjects progressed from one set size to another after a fixed amount of time. Specifically, they spent 178, 204, 260, 354, 370, and 436 seconds on set sizes 1, 2, …, and 6, respectively. These durations were chosen to match the mean duration that subjects in the *E*_0_=45° gamified group spent on the same set sizes. Hence, the trial progression in the control condition roughly matched the trial progression in one of the gamified conditions, but without the presence of any gamified elements. Just as in the gamified conditions, subjects selected at the end of each run whether they wanted to perform 20 extra trials, but without a mentioning of score improvement.

### 2.5. Procedure

Subjects completed the entire experiment in a single session of approximately 90 minutes. After receiving general information about the experiment and signing an informed consent form, they received specific instructions about the task. Subjects in the gamified conditions were told that they would play two rounds of a memory game with the goal to proceed to a level as high as possible. Subjects in the control group were only told that they were going to perform two runs of a memory task with set sizes 1 to 6. Thereafter, the subject performed five practice trials. Control subjects performed the practice trials at set size 1, while subjects in the gamified group performed the trials starting at level = 2.94, to demonstrate the concept of earning points and leveling up. After completing the practice trials, the experiment leader would leave the room and the subject would start the first round of the experiment. After the first round, there was a short break and the subject would start the next round without intervention of the experiment leader. After finishing the second round, the experiment leader would return to the room and conduct a questionnaire, which we describe next.

### 2.6. Questionnaire

We designed a custom questionnaire to obtain insight into aspects related to a subject’s motivation (a copy of it can be found at https://osf.io/gb2kd/). The first part consisted of items similar to the ones found in the Intrinsic Motivation Inventory (IMI; McAuley, Duncan, & Tammen, 1989; Ryan, 1982), such as “I found it interesting” and “It was important for me to perform well”. These items measured motivation in four categories: Interest (items 1, 3, 6, and 8), Perceived Competence (items 5 and 9), Pressure/Tension (items 4 and 10), and Effort/Importance (items 2, 7, and 11). All items were rated on a 1 to 7 integer scale. The second part of the questionnaire consisted of items probing the subject’s mood (“bored”, “frustrated”, etc.) in relation to different set sizes. On hindsight we found no use for the data from the second part and did not include them in the analyses.

### 2.7. Analysis methods

We analyzed the data using Bayesian statistics (Etz & Vandekerckhove, 2018; Wagenmakers, Love, & Marsman, 2018; Wagenmakers, Marsman, et al., 2018). All tests were performed using the JASP software package (JASP Team, 2019) with default prior settings. The subscripts of the Bayes Factors that we report indicate which test was used: BF_10_ indicates the probability of the data under the alternative hypothesis relative to the probability of the data under the null hypothesis; BF_+0_ indicates the probability of the data under the hypothesis that group 1 has a larger mean than group 2, relative to the probability of the data under the null hypothesis; BF_incl_ indicates the probability of the data under models that includes a main effect of the specified factor relative to the probability of the data under models that do not include this main effect. We use the scale provided in Table 1 of Wagenmakers, Love et al. (2018) for interpretation of the strength of evidence (“weak”, “moderate”, etc).

### 2.8. Results

#### 2.8.1. Analysis of motivation scores

We first assess whether gamification affected self-reported scores in the motivation categories Interest, Perceived Competence, Pressure/Tension, and Effort/Importance. For each subject, we compute a single score for each category by averaging across all items within that category. We find that on average, subjects in the gamified conditions reported higher scores in all categories than control subjects (Figure 2). Bayesian t-tests reveal extremely strong evidence for a difference in the category of Interest (BF_+0_=172), strong evidence in the category of Perceived Competence (BF_+0_=88.8), and moderate evidence in the category of Effort/Importance (BF_+0_=3.19). In the category of Pressure/Tension, there was weak evidence in favor of the null hypothesis (BF_10_=0.32). Based on Bayesian ANOVAs with task difficulty as a fixed factor and subject as a random factor, we find for none of the motivation categories evidence that task difficulty affects the self-reported motivation scores (Interest: BF_10_=0.13; Perceived Competence: BF_10_=0.28; Pressure/Tension: BF_10_=0.64; Effort/Importance: BF_10_=0.18). In summary, these data suggest that subjects in the gamified conditions found the task more interesting, felt more competent, and possibly put more effort into it than control subjects.

**Figure 2.**
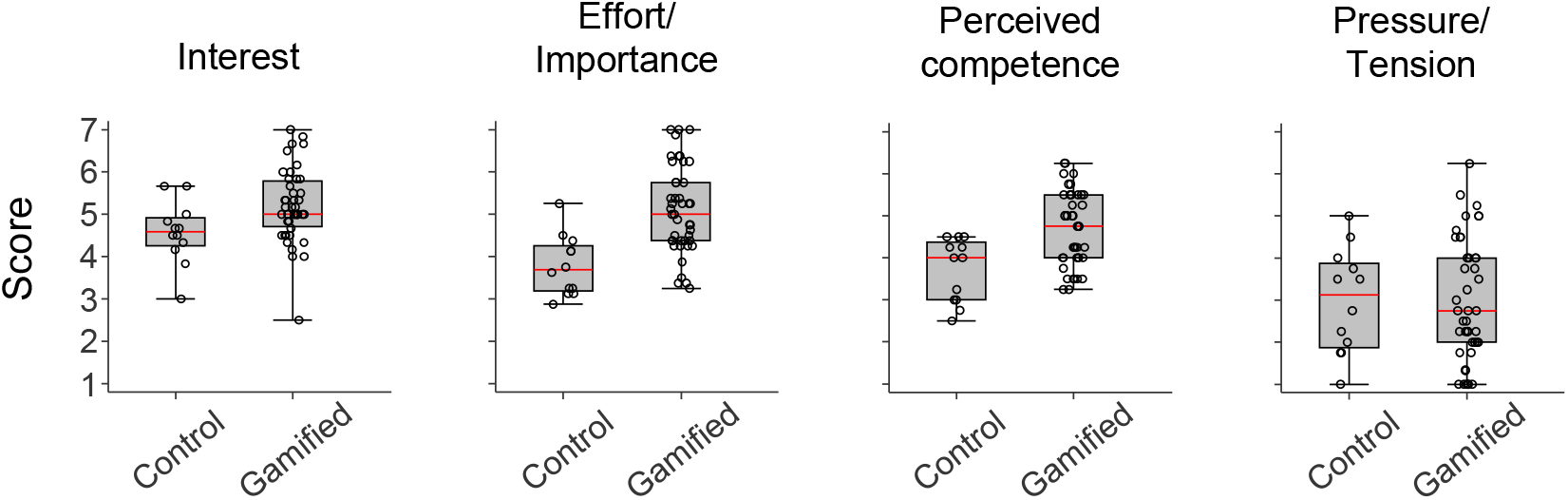
Motivation scores in Experiment 1. Average self-reported scores on the questionnaire, split by motivation category. The boxes indicate the 25% and 75% quantiles, the red line the median, and the whiskers the most extreme values. The circles indicate scores of individual subjects.

#### 2.8.2. Effect of task difficulty on VWM performance

Before we compare performance in the gamified conditions with performance in the control condition, we examine whether there is a difference in average performance between the four gamified conditions. To this end, we perform a Bayesian repeated-measures ANOVA with condition number as a between-subject factor, set size as a repeated measure, and the circular variance of the estimation error as the dependent variable. To avoid the results being affected by “survivor bias”, we restrict all analyses of performance to set sizes for which we have at least 15 measurements for each subject, i.e., set sizes 1 to 4 (at higher set sizes, we lack data for at least one subject in the more difficult conditions). The result of this test provides moderate evidence for the null hypothesis that there is no difference in average performance between the gamified conditions (BF_incl_=0.22). Since both the motivation scores and VWM performance seem unaffected by task difficulty, we treat the four gamified conditions as a single group in the remaining analyses.

#### 2.8.3. Effect of self-reported motivation on VWM performance

We next assess whether the motivation differences between the gamified conditions and the control condition are accompanied by differences in VWM performance. The error histograms suggest that this is not the case, because they look virtually identical between subjects in the gamified conditions and subjects in the control condition (Figure 3A). Indeed, a Bayesian repeated-measures ANOVA with set size as a repeated measure, gamification as a binary between-subjects factor, and the circular variance of the error as the dependent measure reveals moderate evidence in favor of the null hypothesis of there being no effect of gamification on VWM performance (BF_incl_=0.29).

**Figure 3.**
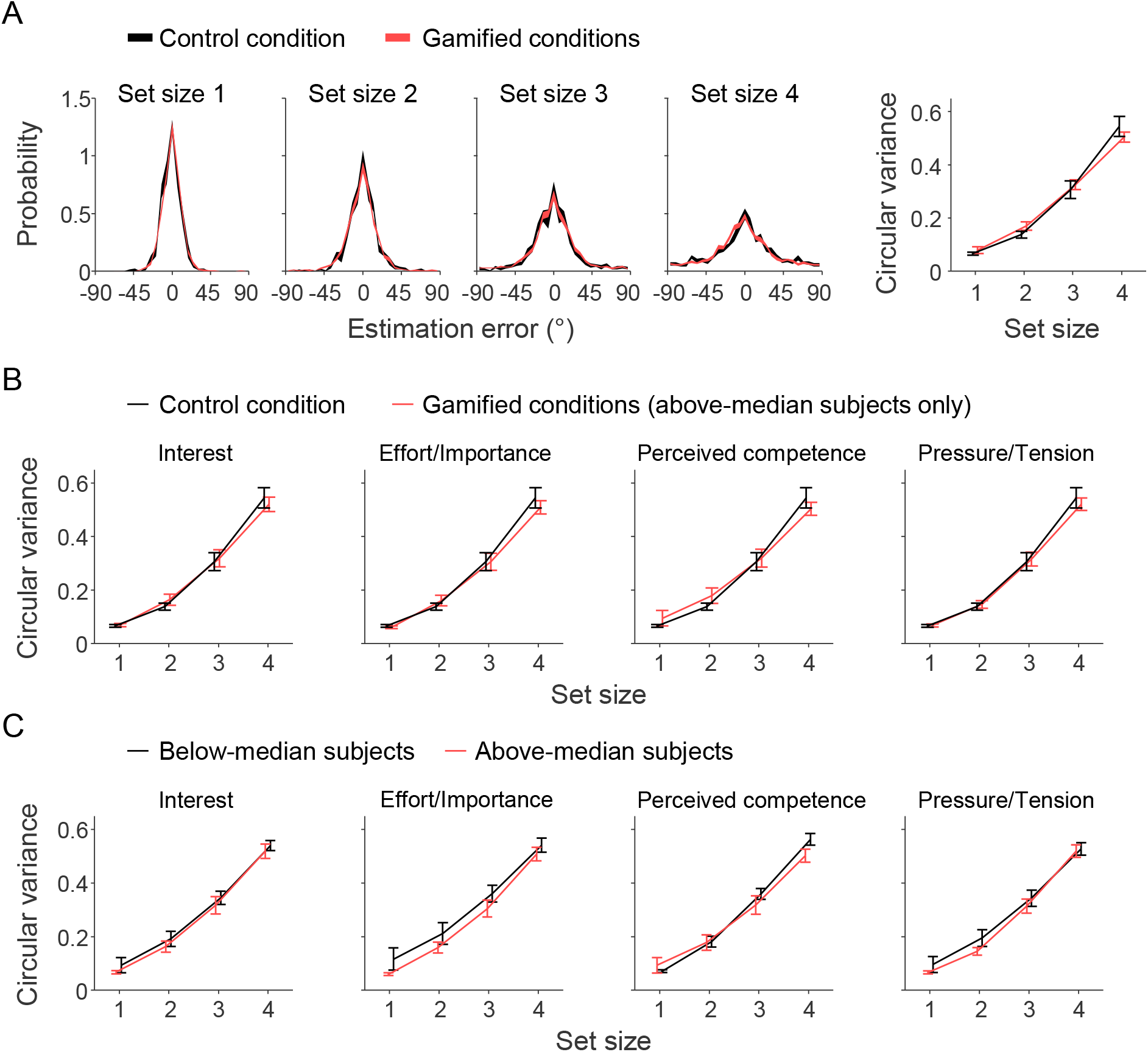
VWM performance in Experiment 1. (A) Estimation error distributions averaged across subjects in the control condition (black) and subjects in the gamified conditions (red). The width the curve indicates one standard error. The graph on the right summarizes the histograms by their circular variance. (B) Circular variance compared between subjects in the control condition and subjects with above-median motivation in the gamified conditions. The median split was performed separately for each motivation category. (C) Circular variance compared between below-medi and above-median subjects in the gamified conditions.

One possible weakness of the previous analysis is that many subjects in the gamified group had motivation scores similar to the control subjects (Figure 2), which suggests that gamification increased motivation for only part of the subjects. Therefore, a more powerful analysis may be achieved by comparing control subjects with only the highly motivated subjects in the gamified conditions. To this end, we divide subjects in the gamified conditions into below and above-median groups based on the motivation scores. We find no evidence that subjects with an above-median Interest score performed on average differently from the control subjects (BF_incl_=0.30). We neither find evidence for an effect when dividing subjects based on a median split in any of the other three motivation categories (Effort/Importance: BF_incl_=0.28; Pressure/Tension: BF_incl_=0.27; Perceived Competence: BF_incl_=0.31). Finally, we compare the above-median subjects in the gamified conditions with below-median subjects (Figure 3C). Again, regardless of which motivation category we use to make the median split, we find no evidence for a difference in VWM performance between the two groups (Interest: BF_incl_=0.26; Effort/Importance: BF_incl_=0.37; Pressure/Tension: BF_incl_=0.26; Perceived Competence: BF_incl_=1.06).

#### 2.8.4. Effect of free-choice behavior on VWM performance

We next analyze whether subjects who voluntarily chose to perform additional trials at the end of a round performed better than subjects who chose not to do so. A total of 41 subjects chose to decline the option in both rounds, 11 subjects performed one extra block of trials, and 10 subjects performed both blocks. We perform a Bayesian repeated-measures ANOVA with set size as a within-subject factor, the number of extra blocks that the subject performed as a between-subject factor, and circular variance as the dependent variable. The result reveals moderate evidence against an effect of the number of extra block (BF_incl_=0.12). Hence, even if subjects who voluntarily chose to perform additional trials were more motivated, this was not accompanied by an increase in performance on the VWM task.

#### 2.8.5. Effect of level progress on VWM performance

So far, we have examined whether there are between-subject differences in performance based on several motivation measures. Next, we assess whether there are any within-subject effects, based on how far a subject had progressed in a level. It could be, for example, that subjects try extra hard when they are close to leveling up or down. For this analysis, we divide the data of subjects in the gamified conditions into 5 bins, with the first bin containing all trials during which the level progress bar was 0-20% filled, the second bin containing all trials during which the bar was filled 20-40%, etc. As before, we only include data from set sizes 1 to 4. A Bayesian ANOVA with bin as a within-subject factor reveals very strong evidence for the null hypothesis of there not being an effect (BF_incl_=0.030).

#### 2.8.6. Effect of round on VWM performance and motivation

Finally, we examine whether there are any differences in motivation or performance between the two experiment rounds. In the control group, scores on Interest, Effort/Importance, and Perceived Competence dropped by 1.31±0.23, 1.31±0.20, 1.00±0.26 points, respectively. The score on Pressure/Tension was similar in both rounds (Figure 4A, left). Consistent with the visual impression, Bayesian paired t-tests provide evidence for a drop in the first three categories (BF_+0_=432, 1.08∙10^3^, and 33.7, respectively), but not in the last one (BF_+0_=0.23). To test whether the differences in motivation scores is accompanied by a difference in VWM performance, we perform a two-way Bayesian ANOVA with set size and round number as independent variables and circular variance as the dependent variable. We find no evidence for a difference of round number on VWM performance (BF_incl_=0.83; Fig 4B, left). Indeed, averaged across all control subjects and set sizes, the relative difference in circular variance between the two rounds is nearly zero (they performed 1.1%±7.2% better in the second round).

**Figure 4.**
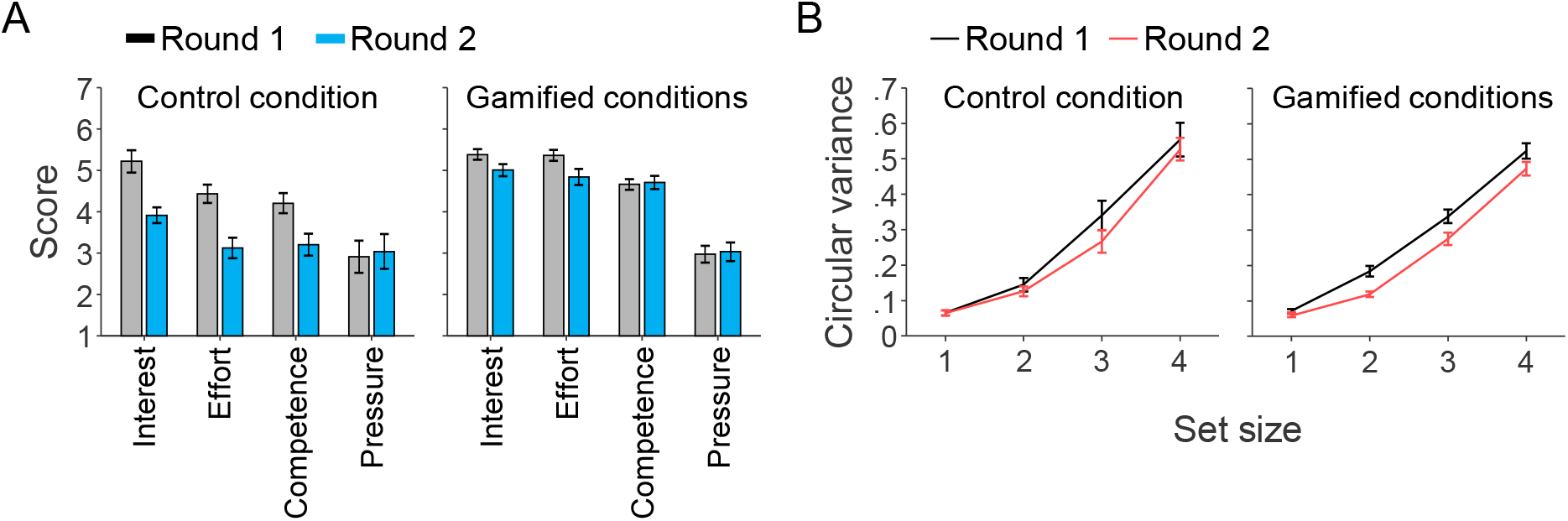
Motivation and performance differences between rounds in Experiment 1. (A) Difference in self-reported motivation scores between the two rounds. (B) Difference in circular variance of the estimation error distribution between the two rounds.

In the group of subjects who performed the gamified version of the experiment, there was no evidence for a drop in Perceived Competence (BF_+0_=0.14) or Pressure/Tension (BF_+0_=0.12) between the first and second round. However, consistent with the control group, there was evidence for a drop in both Interest and Effort/Importance (BF_+0_=78.6 and 48.1, respectively), albeit with much smaller magnitudes (Interest: 0.55±0.15; Effort/Importance: 0.44±0.13; Figure 4B). Interestingly, in contrast to the control group, we find strong evidence for an effect of round on VWM performance in the gamified group (BF_incl_=2.35×10^5^). Further analysis reveals that the circular variance of the error was 8.5±4.3% lower in the second round compared to the first round, indicating an improvement in performance (Figure 4B, right). Since this effect is opposite in direction from what one would expect as a result of the drops observed in Interest and Effort/Importance, we believe that it is best interpreted as a learning effect.

In summary, the results of the comparison between rounds suggest that gamification helps to keep subjects more interested and engaged in the task over a longer period of time. Moreover, we find indications of a learning effect in the gamified conditions, but not in the control condition. One potential explanation is that the sustained engagement of subjects in the gamified tasks was beneficial for learning. However, the lack of a learning effect in the control condition may just as well have been due to lower statistical power (only 12 subjects compared to 50 in the gamified conditions). Most importantly, consistent with the previous analyses, we find no evidence that higher motivation is accompanied by better VWM performance.

### 2.9. Discussion

We gamified a standard VWM task and measured how this affected people’s motivation and performance. While the questionnaire data suggest that subjects in gamified conditions had higher motivation than subjects in the control condition, we did not find any evidence for a difference in memory performance. This null effect on performance is consistent with earlier indications that VWM performance is not sensitive to monetary incentives (van den Berg, Zou, & Ma, 2019; Zhang & Luck, 2011).

The perhaps most straightforward explanation for the absence of an effect is that the total amount of invested VWM resource may be fixed, meaning that no matter how hard subjects try, they cannot improve their performance. This would be somewhat surprising, however, because such inflexibility stands in stark contrast to the flexibility that has been found in how subjects distribute VWM resources across a given set of items: when certain items have higher associated reward (Morey, Cowan, Morey, & Rouder, 2011) or are more likely to be probed than others (Bays, 2014; Emrich et al., 2017; Yoo, Klyszejko, Curtis, & Ma, 2018), subjects assign more resources to that item compared to the other ones. Moreover, it has been found that subjects can increase performance on cued items without a cost for uncued items (Myers, Chekroud, Stokes, & Nobre, 2018) and that specific forms of feedback can improve overall performance on VWM tasks (Adam & Vogel, 2016). Both those findings suggest that it is possible to induce a net increase in utilized VWM resources through experimental manipulations. Also, investing a fixed amount of total resource regardless of the task is suboptimal from a resource-rationality perspective (Van den Berg & Ma, 2018).

An alternative explanation of the null effect is that our experimental design might not have been suitable for inducing or detecting motivation-related flexibility in VWM resource investment. One potential problem is that we used very short stimulus times (200 milliseconds), which may have led to incomplete encoding of the items (Bays et al., 2009). As a result, the maximum precision in VWM was possibly limited by the quality of the input rather than by the amount of available VWM resources. Moreover, it is possible that motivation manipulations are only effective when they are administered on a trial-by-trial basis, as suggested by an earlier study on the relation between task preparation and reward (Shen & Chun, 2011). To address these potential weaknesses in the experimental design, we perform a second experiment with a longer stimulus presentation time and a within-subject manipulation of motivation.

## 3. EXPERIMENT 2

### 3.1. Participants

A total of 12 participants with self-reported normal or corrected-to-normal vision were recruited using posters at various campuses of Uppsala University (8 females; age mean±s.e.m. = 27.6±1.39). The study was approved by the Regional Ethical Review Board in Uppsala and conducted according to the Declaration of Helsinki Principles. All participants signed informed consent and received a cinema voucher with a value of approximately $12 for their participation. One of the participants (S6) had earlier participated as a control subject in Experiment 1.

### 3.2. Experiment procedure

The methods of Experiment 2 were identical to those of Experiment 1, except for the following differences. Most importantly, each subject was tested under three different scoring function with varying amounts of maximum reward (Figure 5B). The scoring functions were of the form 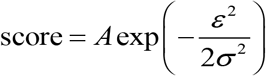 where *ɛ* is the estimation error in degrees, *A* determines the maximum score (obtained when *ɛ*=0), and *σ* determines how quickly the score declines as a function of the error. The “low reward” function always gave 7 points, regardless of the accuracy of the subject’s response (*A*=7, *σ*=∞), the “medium reward” function gave a score between 0 and 21 (*A*=21, *σ*=30), and the “high reward” function gave a score between 0 and 101 (*A*=101, *σ*=20). Subjects were told prior to the experiment that there were trials with low, medium, and high potential reward. However, to avoid that they would challenge themselves to always score the maximum number of points – regardless of the scoring function – we did not tell them the maximum score associated with each function. Within each block of 15 trials, each scoring function was used five times, presented in a random order. To inform the subjects about the reward level of the upcoming trial, we added a reward cue (1.1 sec) at the start of the trial (Figure 5A).

**Figure 5.**
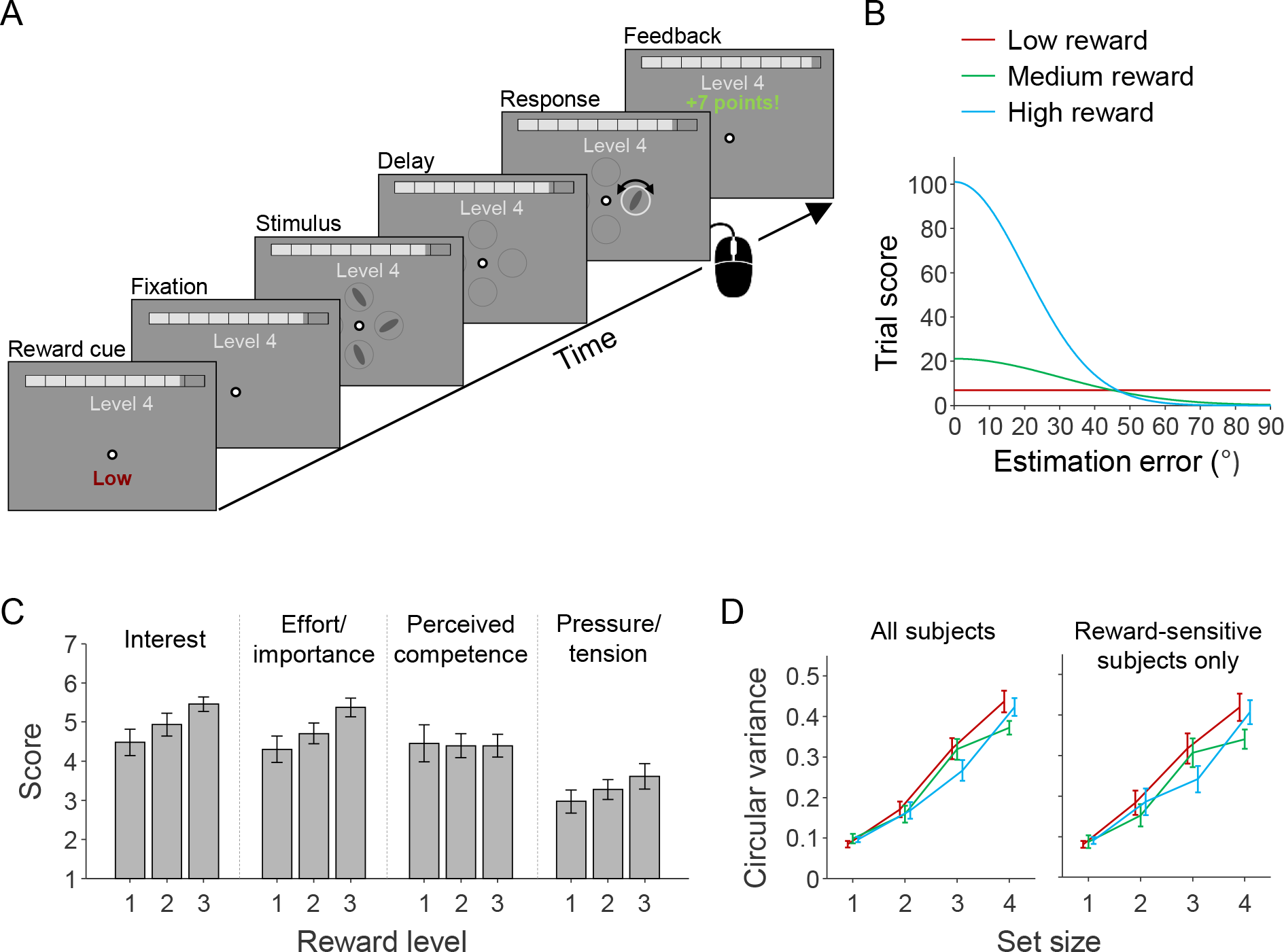
Design and results of Experiment 2. (A) Schematic illustration of a single experimental trial. (B) Scoring functions used in the three trial types. (C) Motivation scores split by category and reward level. (D) Circular variance of the estimation error distribution split by reward level (see panel B for legend), plotted separately for all subjects (left) and for the group of subjects who reported higher motivation for high-reward trials (right).

We made several other minor changes to the experiment. First, we increased the stimulus presentation time from 200 to 900 ms, to reduce the risk that memory quality is limited by the quality of the sensory input rather than the availability of VWM resources. Second, to collect more trials per set size, we increased the number of points required for levelling up from 1,500 to 1,750. Third, we made a minor improvement in how we controlled for fixation errors. In addition to ensuring that subjects were fixating at the start of the trial, we now also forced them to keep fixating during the stimulus presentation and memory delay period. If they broke fixation^3^ during this period, the trial would be terminated with a message “Invalid trial. Please fixate at the center when the stimulus is shown” or “Invalid trial. Please fixate at the center until the response stage”. The time spent on invalid trials was added to the round time, such that each subject spent 30 minutes on valid trials, regardless of the number of invalid trials. Fourth, we added a 30-second forced break between set sizes; after those 30 seconds, subjects could resume the experiment by a keypress whenever they felt ready. Finally, we increased the number of practice trials from 5 to 7 to demonstrate what would happen when breaking of fixation.

### 3.3. Questionnaire

After finishing the experiment, subjects filled out a questionnaire in a web browser. The questions were the same as in the first part of the questionnaire used in Experiment 1, except that a few items were added in each of the four categories (again based on IMI). For all the items we asked separate ratings for low, medium, and high reward trials (a copy of the questionnaire can be found at https://osf.io/gb2kd/). Items 1, 3, 6, 8, 13, and 15 were used to compute Interest scores, items 2, 7, 12, and 17 to compute Effort/Importance scores, items 5, 9, 14, and 16 to compute Perceived Competence scores, and items 4, 10, 11 to compute Pressure/Tension scores. The second part of the questionnaire of Experiment 1 was not included.

### 3.4. Results

A visual inspection of the questionnaire data (Figure 5C) suggests a positive relationship between the reward level (low, medium, high) of a trial and the average motivation score in the categories Interest, Effort/Importance, and Pressure/Tension. To test this statistically, we perform a Bayesian linear regression with Reward Level (coded as 1, 2, or 3) as a covariate and motivation score as the dependent variable. We find moderate evidence for a positive relation in categories Interest (BF_10_=5.03; 95% credible interval on the coefficient: [0.00, 0.46]) and Effort/Importance (BF_10_=3.36, 95% credible interval on the coefficient: [0.00, 0.70]), but not in categories Perceived Competence (BF_10_=0.32) and Pressure/Tension (BF10=0.82). Hence, on average, subjects apparently found the task more interesting at high-reward trials compared to low-reward trials and were willing to put more effort into those trials.

We next examine whether the effect of reward level on motivation was accompanied by an effect on VWM performance (Figure 5D, left). We perform a Bayesian ANOVA with circular variance of the error distribution as the dependent variable, set size and reward level as fixed factors, and subject number as a random factor. Just as in the analyses of Experiment 1, we only include set sizes 1 to 4. The results provide strong evidence for the null hypothesis that there is no effect of reward level on the circular variance (BF_incl_=0.091).

Closer inspection of the questionnaire data reveals that the increase of Interest and Effort/Importance with reward level exists for only 7 of the 12 subjects. The remaining 5 subjects either had a decreasing or non-monotonic pattern in one of the two motivation categories. This indicates that only 7 subjects may have been sensitive to the reward manipulation. To verify that the null result on VWM performance was not due to inclusion of the other 5 subjects in our analysis, we rerun the ANOVA on only the 7 subjects with consistent motivation score patterns (Figure 5D, right). We again find evidence for the null hypothesis that there is no effect of reward level on VWM performance (BF_incl_=0.17).

Another reason for the null result at the group level could be that there is heterogeneity in the effect direction across subjects, such that the effects cancel each other out (e.g., some subjects may have performed better with higher reward, while others might have performed worse). To examine this, we perform a model comparison at the level of individual subjects. The first model is the standard version of the variable-precision model (van den Berg, Awh, & Ma, 2014; van den Berg et al., 2012), which has three free parameters: 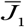 (encoding precision at set size 1), τ (controlling the amount of variability in resource allocation), and *α* (controlling how encoding precision changes with set size). The relation between set size, *N*, and encoding precision, 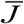, is defined as 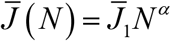. We use 5-fold cross validation to compare the fit of this model with a variant in which encoding precision depends on both set size and reward level *R* (coded as 1, 2, and 3): 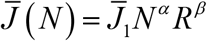, where *β* is an additional free parameter. When *β*=0, the two models are identical. For subjects with a strong relation between reward level and VWM performance, *β*=0 will not provide a good fit. For those subjects we should find a strong advantage of the second model over the first one. By contrast, we find that for every subject the cross-validated log likelihood difference between the two models is close to zero (range: 1.05 in favor of the first model to 1.99 points in favor of the second model). In fact, for 10 of the subjects the first model is favored. These results corroborate the earlier evidence suggesting that VWM performance was unaffected by reward level.

### 3.5. Discussion

The purpose of Experiment 2 was to address two potential weaknesses in Experiment 1 by increasing the stimulus presentation time and manipulating motivation on a trial-by-trial basis. The post-questionnaire data of Experiment 2 showed a positive relation between Interest and Effort/Important scores on the one hand and reward level on the other hand, indicating that our within-subject design had the desired effect on motivation. However, the increase in motivation on high-reward trials was not accompanied by an increase in VWM performance. Hence, the results of Experiment 2 are consistent with those of Experiment 1: gamification had a positive effect on motivation, but left performance unaffected.

## 4. GENERAL DISCUSSION

While it is known that gamification can increase subjects’ intrinsic motivation, enjoyment, engagement, and perceived competence in a task (Hamari et al., 2014), little is known about effects of gamification on cognitive performance. Here, we performed two experiments to investigate whether gamification of a standard VWM task improves subjects’ performance on that task. Consistent with previous literature, we found positive effects of gamification on subjects’ self-reported interest in the task and the amount of effort they reportedly put into it. However, in neither experiment did we find any differences in VWM performance. We thus conclude that gamification can make people more motivated to perform VWM tasks, but it does not necessarily make them better at it.

Our finding that there was no relation whatsoever between motivation and VWM performance in our experiments is puzzling for several reasons. First, there is convincing evidence that people are able to flexibly assign more VWM resources to important items compared to less important ones, in a way that seemingly optimizes performance (Bays, 2014; Emrich et al., 2017; Morey et al., 2011; Yoo et al., 2018). As we have demonstrated earlier (van den Berg & Ma, 2018), it would be suboptimal if people would always invest the same amount of VWM, independent of task properties. Our finding that they nevertheless seemed to do so in our experiments raises the question why VWM has evolved to be approximately optimal in distributing a given amount of resource across items, but at the same time uses a highly suboptimal policy to determine the total amount of VWM resource to invest in a task. Moreover, it is known that motivation and expected reward affect dopamine concentration (e.g., Di Chiara, 2005; Hamid et al., 2015; Syed et al., 2015) and that dopamine concentration, in turn, is related to VWM performance (e.g., Arnsten, 1998; Ashby & Valentin, 2017; Cools et al., 2008; Okimura et al., 2015; Sawaguchi & Goldman-Rakic, 1991; Williams & Goldman-Rakic, 1995). Based on those findings, one would expect – contrary to our findings – that any experimental manipulation that affects participants’ motivation, also affects their VWM performance.

An alternative explanation for our null findings is that subjects may have been overperforming: regardless of the condition they were in, they may have felt a responsibility to perform as well as they could, for instance as a justification for their payment, out of a desire to deliver high-quality data to the researchers, or out of fear to be confronted with their performance after finishing the experiment. Based on informal feedback, we know that at least one of the subjects who voluntarily performed the additional trials did so because she believed it would help the researchers. While most studies that manipulate motivation do so by incentivizing subjects, if overperformance is a real issue, it might be interesting for future studies to look into ways to *decrease* motivation levels. Moreover, it may be that highly artificial stimuli are not very suitable for manipulating motivation levels, for instance due to a lack of engagement in the task. Therefore, it might also be fruitful if future studies would use more naturalistic stimuli. Another potential limitation of the present study is that we used a questionnaire to measure how motivated subjects were and how much effort they put into the task. Another limitation of the present study is that we only analyzed VWM performance at relatively low set sizes (1 to 4). We excluded higher set sizes in order to avoid “survivor bias” effects in the results. However, the subjects’ answers to the questionnaire items that we used to measure intrinsic motivation were based on their experience of all set sizes they performed. Hence, the motivation data may be unreliable for the set sizes that we used to analyze VWM performance differences. To address this problem, it would have been good to complement the self-report measures with pupillary dilation data, which is known to correlate with attention and effort (Hoeks & Levelt, 1993; van der Wel & van Steenbergen, 2018). In the present study we collected eye movement data, including pupil dilation measurements, but we did so mainly to verify that subjects were fixating and not with the aim to use them in our analyses. As a result, the collected data are unfortunately unsuitable for analysis.

As a final remark, we believe that our study may serve as a showcase of open science in a field that is currently plagued by concerns about replicability (Aarts et al., 2015). Although an increasing number of authors make their data available – partly driven by changes in journal policies – researchers still seem to be wary of publishing null results, especially when this a result contradicts their own theory. Keeping such findings hidden in a file drawer may have short-term benefits for the researchers, but is seen as one of the major factors behind the replicability crisis.

## ACKNOWLEDGEMENTS

RvdB is supported by Grant 2018-01947 from the Swedish Research Council (Vetenskapsrådet) and Grant RIK17-1157:2 of the Bank of Sweden Tercentenary Foundation (Stiftelsen Riksbankens Jubileumsfond). The funding sources had no role in the study design, the collection, analysis and interpretation of data, the writing of the report, and the decision to submit the article for publication.

## Footnotes

1 The room contained two ceiling-mounted fluorescent lamps. The one farthest away from the experimental setup was turned on and the other on was turned off. However, a number of subjects reported that the lamp closest to the setup had spontaneously turned on during the experiment, possibly due to a technical error in the lamp’s motion detector. Moreover, one subject was accidentally tested in a completely darkened room.

2 More precisely, the level was computed at the beginning of each trial as 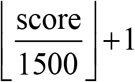, where the 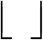 operation rounds a number down to the closest integer

3 We considered fixation to be broken once the eye tracker returned 5 measurements in which the gaze location was farther than 150 pixels away from the center of the fixation dot.

